# Repeated evolution of asexuality involves convergent gene expression changes

**DOI:** 10.1101/364869

**Authors:** Darren J. Parker, Jens Bast, Kirsten Jalvingh, Zoé Dumas, Marc Robinson-Rechavi, Tanja Schwander

## Abstract

Asexual reproduction has evolved repeatedly from sexual ancestors across a wide range of taxa. While the costs and benefits associated with asexuality have received considerable attention, the molecular changes underpinning the evolution of asexual reproduction remain relatively unexplored. In particular, it is completely unknown whether the repeated evolution of asexual phenotypes involves similar molecular changes, as previous studies have focused on changes occurring in single lineages. Here we investigate the extent of convergent gene expression changes across five independent transitions to asexuality in stick insects. We compared gene expression of asexual females to females of close sexual relatives in whole-bodies, reproductive tracts, and legs. We identified a striking amount of convergent gene expression change (up to 8% of genes), greatly exceeding that expected by chance. Convergent changes were also tissue-specific, and most likely driven by selection for functional changes. Genes showing convergent changes in the reproductive tract were associated with meiotic spindle formation and centrosome organization. These genes are particularly interesting as they can influence the production of unreduced eggs, a key barrier to asexual reproduction. Changes in legs and whole-bodies were likely involved in female sexual trait decay, with enrichment in terms such as sperm-storage and pigmentation. By identifying changes occurring across multiple independent transitions to asexuality, our results provide a rare insight into the molecular basis of asexual phenotypes and suggest that the evolutionary path to asexuality is highly constrained, requiring repeated changes to the same key genes.

---

Sexual reproduction is extremely costly. Sex is less efficient than asexuality for transmitting genes to future generations (Maynard Smith 1971) and in order to outcross, an individual has to find a partner, forgo foraging, and risk contracting sexually transmitted diseases and predation while mating (Bell 1982; Lehtonen et al. 2012). Yet, the overwhelming number of sexual, as compared to asexual, animal and plant species (Avise 2008; van der Kooi et al. 2017) indicates that sexual reproduction is highly advantageous. Identifying potential advantages conferred by sex has motivated decades of research and a rich body of work on the evolution and maintenance of sexual and asexual reproduction has been produced (reviewed in Bell 1982; Lewis 1987; West et al. 1999; Otto 2009; Neiman et al. 2017). By contrast, little is known about the molecular underpinnings required to evolve asexuality from sexual ancestors (Neiman et al. 2014). Yet these molecular underpinnings have the potential to provide insights into the processes involved in the evolution of asexuality, and to help understand how sex is maintained. For example, sex is more easily maintained if asexuality evolves gradually in a sexual population than if it emerges suddenly via major effect mutations (Templeton 1982; Burt 2000; Schwander et al. 2010).

Some insight into the genetic basis of asexuality has been gained from studies of individual asexual lineages (Innes and Hebert 1988; Lynch et al. 2008; Eads et al. 2012; Jaquiéry et al. 2014), but a broad comparative framework for exploring common principles of the molecular basis of asexuality is lacking. For example, a major unresolved question is whether independent transitions to asexuality involve similar or different molecular changes. To address these shortcomings, we explored the molecular underpinnings of asexuality in stick insects of the genus *Timema*, a genus of wingless, herbivorous insects native to the West coast of North America and the mountains of the Desert Southwest. This group is uniquely suited for comparative studies of asexuality, as asexuality has evolved at least seven times independently (Schwander et al. 2011) (Fig. 1), allowing us to study convergence across replicate transitions from sexual to asexual reproduction. Furthermore, close sexual relatives are at hand for each asexual lineage for comparison. All asexual *Timema* species reproduce via obligate parthenogenesis (Schwander and Crespi 2009), meaning that they evolved the ability to produce unreduced eggs which develop without fertilization by sperm. Additional phenotypic changes evolved convergently as adaptations to asexuality, including a reduced sperm storage organ, and reduced sexual pheromone production (Schwander, Crespi, et al. 2013). Thus, asexual *Timema* females are less attractive to sexual males (Schwander, Crespi, et al. 2013), which use both airborne and contact signals to identify suitable mates (Nosil et al. 2007; Arbuthnott and Crespi 2009; Schwander, Arbuthnott, et al. 2013), and even when copulations between sexual asexual females and males from sister-species are forced under laboratory conditions, eggs are not fertilized (Schwander, Crespi, et al. 2013).

**Figure 1.**
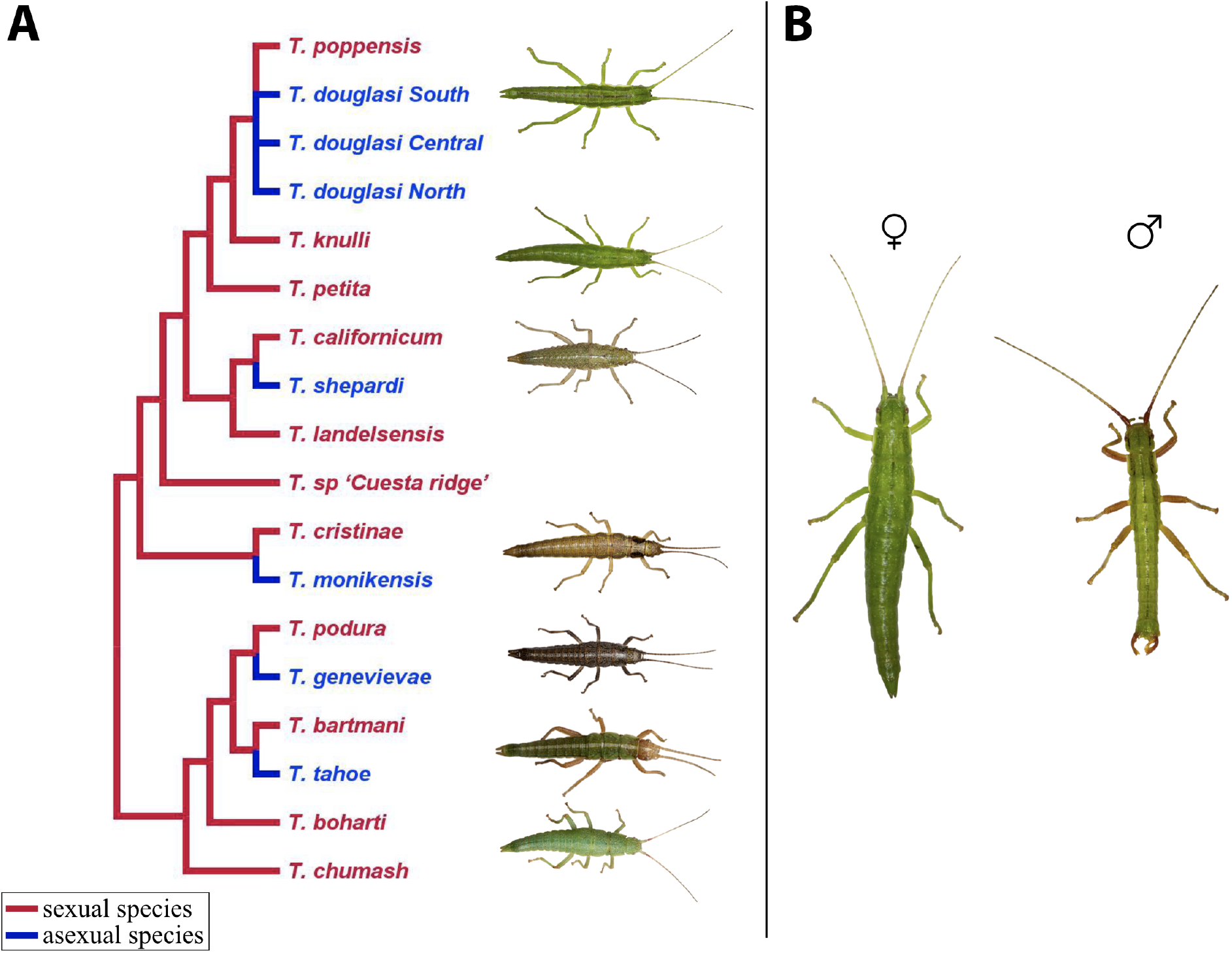
**A.** Phylogeny of described *Timema* species (redrawn from (2017) with asexual species added from (2011)). Sexually reproducing species are shown in red, independently derived asexual lineages in blue. Our study used the five asexual species (for *T. douglasi* only the southern lineage was used (labelled as *T. douglasi* South)) and their sexual sister species. **B.** Sexual dimorphism in *Timema* (*T. knulli*).

To capture molecular changes associated with the evolution of asexuality we performed whole-body and tissue-specific transcriptome sequencing (RNA-seq) on females from five sexual and five asexual *Timema* species (Fig. 1). We chose two different tissues, the reproductive tract and legs, to identify the molecular mechanisms underlying the production of asexual offspring (reproductive tract), and adaptations to a celibate life (e.g. reduction of various different sexual traits in the reproductive tract and legs). Note that the reproductive tract and leg samples actually represent a collection of tissues, but we refer to them as tissues throughout for brevity. Whole-body samples were included as they allow us to identify important changes that may be missing in the tissue-specific transcriptomes. Using this approach, we identified convergent expression changes which were likely driven by selection. We also observed changes specific to each sexual-asexual species-pair which typically showed concerted changes across tissues, consistent with being a product of drift (Blekhman et al. 2008; Liang et al. 2018). Finally, to complement our expression analyses, we examined patterns of molecular evolution in genes showing convergent expression changes following a transition to asexuality.

## Results

### Transcriptomes and orthology

Reference transcriptome assemblies for each species were generated previously (Bast et al. 2018). Bast et al. (2018) also identified 3010 one-to-one orthologs, which were used as our transcriptome reference. For each tissue, orthologs with low expression (counts per million less than 0.5 in two or more libraries per species) were filtered prior to expression analyses. Thus, the final number of orthologs kept for analyses of whole-body, reproductive tract, and leg samples was 2984, 2753, and 2740, respectively.

### Convergent gene expression changes

We identified convergent gene expression changes between sexual and asexual species by modelling gene expression as a function of species-pair (see Fig. 1), reproductive mode (sexual or asexual), and their interaction in edgeR (Robinson et al. 2010). In such a model, convergence is indicated by an overall effect of reproductive mode (FDR < 0.05), but no interaction (FDR > 0.05) (Supplemental Table 1). Approximately four times as many genes changed convergently in the reproductive tract (7%; 203/2754) and legs (8%; 206/2737) as compared to the whole-body (2%; 57/2985), perhaps reflecting the relative difficulty in identifying expression changes in complex tissue assemblies such as whole-bodies (Johnson et al. 2013). The amount of convergence we observe is considerable and approximately double what we would expect by chance, for all tissues (whole-body: p = 0.0128, reproductive tract: p < 0.0001, legs: p < 0.0001, Supplemental Fig. 1). The amount of change between sexual and asexual females was relatively small for convergent genes, with a mean fold change of approximately 1.4 (absolute log_2_ expression change for whole-body = 0.55, reproductive tract = 0.68, and legs = 0.46) (Fig. 2).

**Figure 2.**
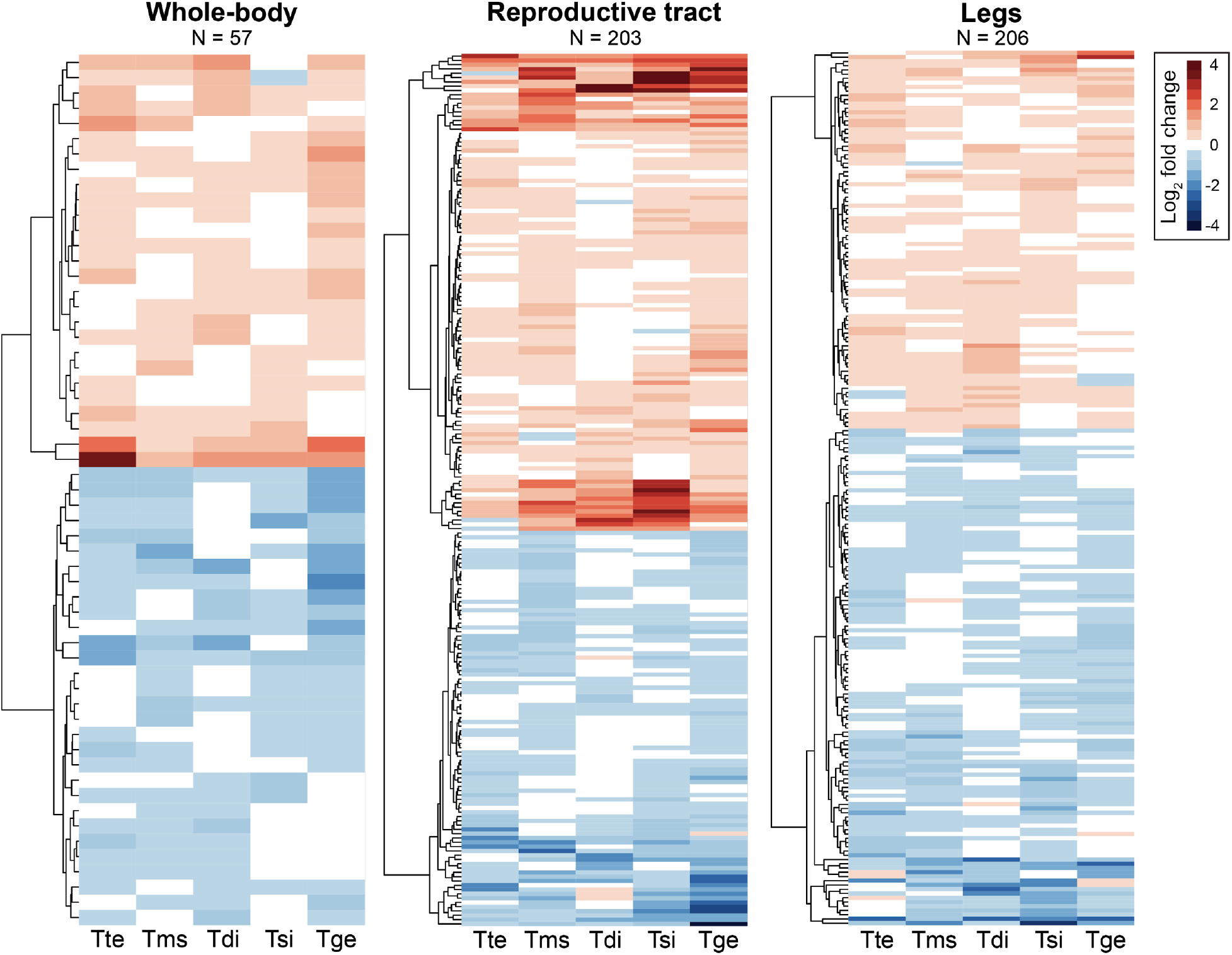
Heatmaps of genes showing convergent gene expression changes between sexual and asexual females for whole-bodies, reproductive tract, and legs. Species names are abbreviated as follows: Tte = *T. tahoe*, Tms = *T. monikensis*, Tdi = *T. douglasi*, Tsi = *T. shepardi*, and Tge = *T. genevievae*.

Genetic changes that influence gene expression are likely to cause correlated changes in expression across tissues (GTEx Consortium. 2017; Liang et al. 2018). If such changes are non-neutral they are likely to be deleterious due to pleiotropy. As a consequence, gene expression changes that occur in parallel across different tissues are more likely a result of drift rather than selection (Blekhman et al. 2008; Liang et al. 2018). We found that convergent changes between sexual and asexual species were highly tissue-specific. Only 22 of the convergent genes in the reproductive tract (203) and legs (206) overlapped between the two tissues, a value not significantly greater than expected by chance (Table 1). There was also little overlap between convergent genes in the two tissues and whole-bodies (Table 1, Supplemental Fig. 2). This pattern is consistent with the idea that convergent expression changes in asexuals are driven by selection rather than by drift. This interpretation of selection also predicts that convergent genes will be involved in divergent functions in each tissue which, as we show below, is indeed the case.

**Table 1.** Overlap of differentially expressed genes between different tissue types. Number of genes expected by chance given in parentheses. P-values are from a fisher’s exact test corrected for multiple tests. Species names are abbreviated as follows: Tbi = *T. bartmani*, Tce =*T. cristinae*, Tps = *T. poppensis*, Tcm = *T. californicum*, Tpa = *T. podura*, Tte = *T. tahoe*, Tms = *T. monikensis*, Tdi = *T. douglasi*, Tsi = *T. shepardi*, and Tge = *T. genevievae*.

|  | Reproductive tract & legs | Whole-body & legs | Whole-body & reproductive tract |
| --- | --- | --- | --- |
| **Convergent genes** | 22 (16)<br>$p = 0.14$ | 7 (5)<br>$p = 0.21$ | 4 (4)<br>$p = 0.65$ |
| **Species-pair (Tbi-Tte)** | 62 (24)<br>$p = 1.30 \times 10^{-14}$ | 68 (29)<br>$p = 4.57 \times 10^{-14}$ | 25 (7)<br>$p = 4.53 \times 10^{-9}$ |
| **Species-pair (Tce-Tms)** | 88 (47)<br>$p = 1.36 \times 10^{-10}$ | 48 (9)<br>$p = 6.19 \times 10^{-25}$ | 47 (14)<br>$p = 3.86 \times 10^{-16}$ |
| **Species-pair (Tcm-Tsi)** | 42 (11)<br>$p = 4.45 \times 10^{-15}$ | 24 (5)<br>$p = 4.64 \times 10^{-11}$ | 19 (3)<br>$p = 4.64 \times 10^{-11}$ |
| **Species-pair (Tpa-Tge)** | 153 (134)<br>$p = 1.99 \times 10^{-2}$ | 102 (60)<br>$p = 1.22 \times 10^{-9}$ | 86 (54)<br>$p = 8.75 \times 10^{-07}$ |
| **Species-pair (Tps-Tdi)** | 56 (20)<br>$p = 1.31 \times 10^{-15}$ | 59 (37)<br>$p = 6.08 \times 10^{-5}$ | 32 (7)<br>$p = 7.81 \times 10^{-14}$ |

To directly test whether the convergent changes are driven by selection, we modeled convergent change as an Ornstein-Uhlenbeck (OU) process. Selection-driven expression changes are identified in this framework by comparing the likelihood of two models: a drift-model where expression changes are modelled as simple Brownian motion, and an adaptive-optima model where expression changes are modelled as the result of both Brownian motion and selection toward specified adaptive-optima (see Butler and King (2004) and Cressler et al. (2015)). In this case the second model specified that asexual species had a different adaptive-optimum than sexual species. As each asexual species is phylogenetically independent, this tests for convergence of expression. Using this approach, we found that the adaptive-optima model had a significantly better fit than the drift model for 70% to 80% of convergently expressed genes, depending on tissue (Supplemental Table 2 and 3, FDR < 0.05). This strengthens our interpretation that the majority of the convergent changes between sexual and asexual species are driven by selection.

Finally, we compared the two-state adaptive-optima model, where all asexual species share the same optimum, to a model where the optimum can vary between asexual species (multi-asexual optima model). We find that approximately 10% of the convergently expressed genes have a better fit to the multi-asexual optima model than to the two-state model (Supplemental Table 2 and 3), showing that for some genes, the convergent increase or decrease in gene expression differs in magnitude in different asexual species (Supplemental Fig. 3).

### Functional processes of convergently expressed genes

To detect convergence at the process level, we performed gene set enrichment analyses (GSEA) for each tissue separately. Briefly, we scored Gene Ontology (GO) terms according to the rank of convergent expression change of genes annotated to the terms; GO terms were then called significant if they had a better average rank than expected by chance (see Methods). For each tissue, more than 100 GO terms are enriched (FDR < 0.05), providing strong support for convergence of biological processes between asexual species (Supplementary Tables 4-6). This signal is not dependent on any threshold at the gene level, and thus provides information on convergence at the process level due to small but consistent contributions from many genes. Consistent with the gene expression results, enriched GO terms were generally tissue specific; we found no significant overlap between GO-terms enriched in the legs and reproductive tract (11 shared terms, FDR = 0.123), between the legs and whole-body (4 shared terms, FDR = 0.799), or between whole-body and reproductive tract samples (10 shared terms, FDR = 0.064).

To reduce the number of enriched GO terms to examine we semantically clustered enriched GO terms using ReviGO (Supek et al. 2011) (Supplemental Tables 7-9). The annotations of convergent changes in the reproductive tract reflect the convergent evolution of parthenogenesis in asexual *Timema*, as they were linked to meiosis (meiotic spindle organization, meiosis II, centrosome duplication, meiosis I cytokinesis, meiosis II cytokinesis), and reproduction (growth of a germarium-derived egg chamber, sperm individualization, gamete generation). However, convergent changes were also linked to neuron development (neurogenesis, neuron development, neuron recognition), as well as several GO terms involved in development and metabolic processes for which the link to asexuality is less clear. In legs we identified GO terms involved in immune defence (response to fungus, regulation of production of molecular mediator of immune response, regulation of antimicrobial peptide production, regulation of humoral immune response), which may be because asexual females are no longer susceptible to the costs associated with diseases transmitted from sexual interactions (Knell and Webberley 2004). Convergent changes were also linked to sex determination (primary sex determination; soma, primary response to X:A ratio), which may control changes in the expression of sexual traits, and several metabolic processes. In whole-body samples we find some reproduction associated terms (courtship behavior, male mating behavior, male courtship behavior, sperm storage, regulation of ovulation) as in the reproductive tract, and behavioral, and immune related terms (immune response-regulating cell surface receptor signaling pathway) as in legs, but also some unique terms relating to the cuticle (ecdysone, pupal chitin-based cuticle development).

### Convergently expressed genes in whole-bodies show evidence for sexual trait decay

Several of the enriched functional processes described above are suggestive of sexual trait decay. Under this scenario we expect a reduction of purifying selection on genes underlying sexually dimorphic traits in asexual species, indicated by an increased accumulation of non-synonymous changes.

The power to detect differences in pN/pS or dN/dS between gene sets in asexuals is low, as genes are inherited as a single linkage group. Nevertheless, we found that genes showing convergent changes in expression in whole-bodies showed elevated pN/pS and dN/dS when compared to the genomic background (permuted t-test p-value for: pN/pS < 0.0001, dN/dS = 0.0084, Fig. 3, Supplemental Figs. 4 and 5), consistent with the idea of sexual trait decay. Sexual trait decay is further supported by the examination of functional annotations for such genes which include one gene (OG-2854) that is produced primarily in male accessory glands in *Drosophila*, and at least three other genes (OG-2197, OG-663, OG-1014) that are involved in pigment synthesis pathways (pigmentation is sexually dimorphic in *Timema* see Fig. 1B). In contrast, genes showing convergent changes in expression in the reproductive tract and legs did not show elevated pN/pS or dN/dS (Fig. 3, Supplemental Figs. 4 and 5), suggesting that convergent expression changes in these tissues do not coincide with reduced purifying selection acting on their sequences.

**Figure 3.**
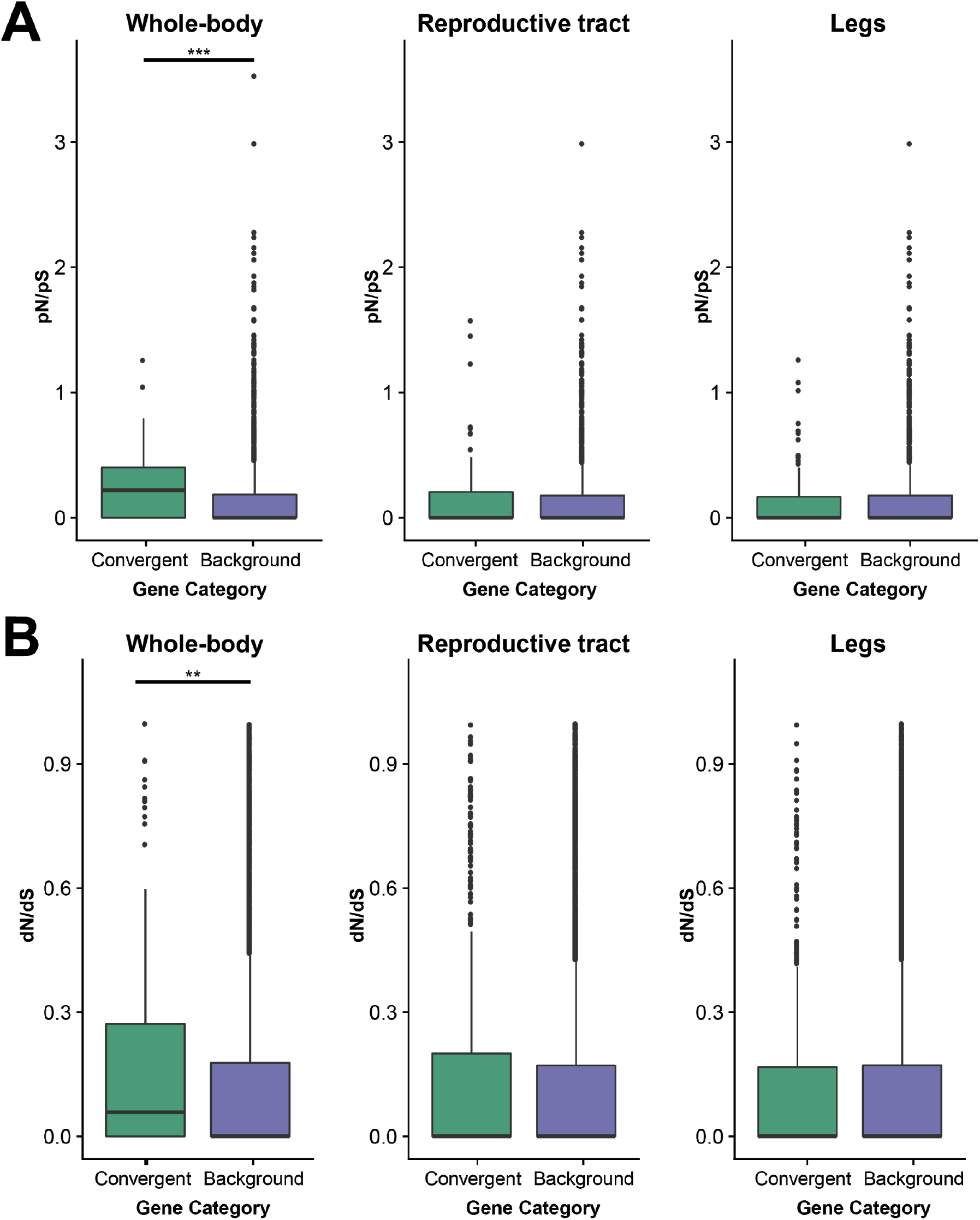
pN/pS ratios (A) and dN/dS ratios (B) for convergently expressed genes versus all other genes expressed in that tissue for whole-bodies, reproductive tracts and legs. Significance is indicated by asterisks (** < 0.01, *** < 0.001).

We conducted several additional analyses to check the robustness of our results and corroborate our interpretations. Firstly, we examined in detail the associated functional annotations of candidate gene sets for which there was very strong evidence for convergent changes, and secondly, we used cross-species mapping to examine expression changes occurring across the whole transcriptome, rather than only in the subset of genes we identified as single copy orthologs between the 10 species. Both approaches support the results from our original analyses and are described below.

### Strongly convergent candidate genes and their function

Although all the convergent genes we identified showed an overall shift in expression across the five species-pairs, often expression change in one or two of the pairs was small (<1.2 fold change). We defined top candidate genes as convergent genes for which the absolute log_2_ fold change in expression was more than 0.25 (~1.2 fold change) for all species-pairs. Most of these top genes showed convergent shifts in the reproductive tract (36 genes, relative to 4 and 15 genes for whole-body and legs, respectively) (Figure 4, Supplemental Table 1). The functions of these candidate genes largely reflected the functional processes identified for the full set of convergently expressed genes, and highlight a number of key genes potentially involved in producing asexual phenotypes.

**Figure 4.**
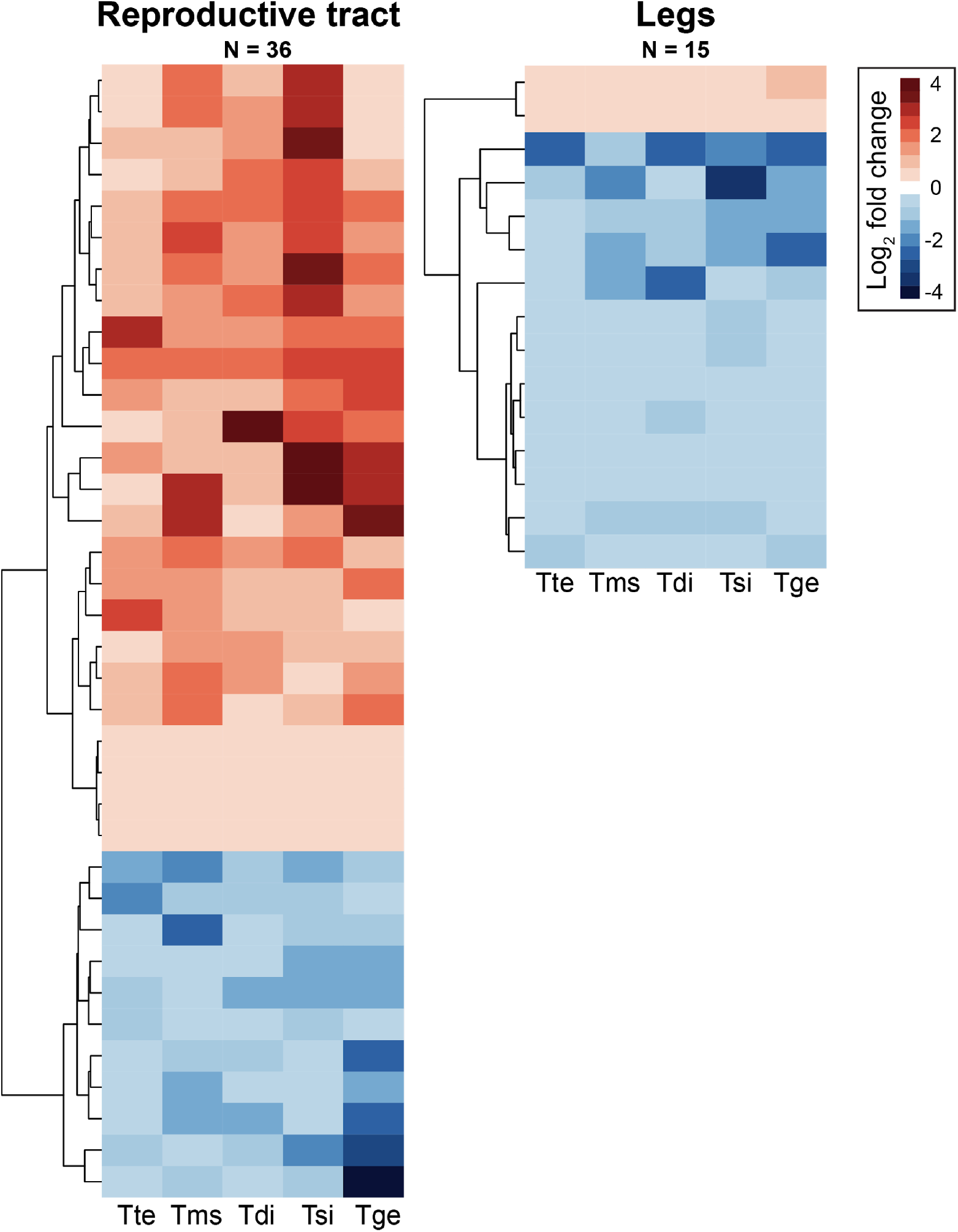
Heatmaps of candidate convergent gene expression changes between sexual and asexual females for the reproductive tract and legs. Species names are abbreviated as follows: Tte = *T. tahoe*, Tms = *T. monikensis*, Tdi = *T. douglasi*, Tsi = *T. shepardi*, and Tge = *T. genevievae*.

For the reproductive tract four genes are involved in meiotic spindle formation and centrosome organization (OG-513, OG-1448, OG-1488, OG-314). In particular we find two genes (OG-1448, OG-1488) belonging to a family of Elovl proteins that mediate elongation of very-long-chain fatty acids, including an ortholog to *D. melanogaster* gene *bond*, which effects spindle formation and has been shown to be important for meiotic, but not mitotic, cytokinesis (Szafer-Glusman et al. 2008). In particular, *D. melanogaster* males defective for *bond* commonly display two to four nuclei in spermatids causing sterility. Female *bond* mutants are also infertile (Szafer-Glusman et al. 2008), although the mechanism is unknown. The other two genes (OG-513, OG-314) have roles in centrosome function, including an ortholog to *poc1* which is involved in centrosome formation (Blachon et al. 2009). Six genes (OG-758, OG-2002, OG-1478, OG-1993, OG-2686, OG-148) were annotated with reproduction associated terms which may be responsible for the convergent reproductive changes we observe between asexual and sexual females. Interestingly, one gene, OG-511, is an ortholog to glucose dehydrogenase which is important for sperm storage in female *D. melanogaster* (Bloch Qazi et al. 2003). Finally, we find that 11 genes (OG-1195, OG-1478, OG-1841, OG-2197, OG-2808, OG-366, OG-445, OG-511, OG-705, OG-712, OG-758, OG-810) have annotations to the nervous system. The majority of these appear to be sensory in nature, and in particular seven are annotated with the GO term “sensory perception of pain”. Changes in these genes may represent changes associated with female receptivity and post-mating behaviour in asexual females, which are target of substances in the male ejaculate (Heifetz and Wolfner 2004; Sakai et al. 2009; Heifetz et al. 2014).

For leg samples three genes (OG-1651, OG-2048 and OG-1081) are involved in immune defence. In particular orthologs of both genes (*Trx-2* and *MP1*) are involved in the activation of melanisation in response to fungal and bacterial infection in *D. melanogaster* (Tang et al. 2006; Jin et al. 2008). Three genes are involved in cuticle development (OG-2221, OG-2738, and OG-2995). Orthologs of two other genes (OG-1371 and OG-2031) are involved in male specific behaviours (male courtship behavior and inter-male aggressive behavior) in *D. melanogaster* (*CaMKII* and *Fkbp14*) (Mehren and Griffith 2006; Edwards et al. 2009). Since these genes are also expressed in females, changes to their expression may have resulted from the release of intralocus sexual conflict.

Whole-body samples had only four strong candidate genes, and all either have no annotation or have only broad GO-terms annotated. One potentially interesting gene, OG-2188, has an ortholog (*CG12237*) that has been associated with female sterility in *D. melanogaster* (Sopko et al. 2014). Finally, the remaining candidate genes across all tissues were either unannotated (12 genes) or only have very broad GO-terms annotated (10 genes).

### Cross-species mapping

Using only the 3010 genes with 1-to-1 orthologs across all species could impact our ability to detect convergent changes since we only use a relatively small fraction of the total number of transcripts in each assembly (23435 to 37847; Supplemental Table 10). To investigate more genes, we mapped reads from all samples to genes from each species which had a reciprocal-best-blast-hit between sexual-asexual sister species (which includes the 1-to-1 orthologs analysed above). This approach generated 10 different datasets (one for each species assembly), with between 15500 and 17583 genes. After filtering out genes with low expression (using cpm, see Methods) in each dataset, this approach allowed us to examine between 2.43-3.12 (dependent on species and tissue) times more genes than using the 1-to-1 orthologs (Supplemental Table 10). Results from this approach qualitatively confirmed the results found using only the 1-to-1 orthologs: the percentage of genes showing a convergent expression ranged from 4-5% for whole-body samples and 6-8% for the reproductive tract and leg samples, dependent on which of the species transcriptome was used (Supplemental Table 10), and GSEA produced similar enriched GO terms (Supplemental Tables 11-13)

### Species-pair specific changes

The approach taken above allowed us to identify genes which showed convergent changes in expression across independent transitions to asexuality. This approach will not identify expression changes confined to a single or a few species-pairs. Changes occurring in only a minority of species-pairs are clearly not convergent at the gene expression level, however, these changes could be convergent at the functional process level, whereby species-pair specific changes in gene expression are involved in common functional processes between species-pairs (Rittschof et al. 2014; Berens et al. 2015). To test this, we compared each asexual species to its closest sexual relative and called differentially expressed (DE) genes from each pairwise comparison.

The number of significantly DE genes between each pair varied greatly depending on species-pair and tissue, with a generally greater number of genes DE in leg tissue (59 to 626, Supplementary Fig. 6). This greater number in leg tissue is likely due to the smaller variation between replicates (common biological coefficient of variation was lowest for legs: whole-body = 0.314, reproductive tract = 0.340, legs = 0.238), as tissue differences disappeared when a fold-change threshold was applied (Supplementary Fig. 7). There were no genes that showed overlap between all sexual-asexual species-pairs in any tissue (Fig. 5A). Examination of overlaps between pairs of sexual-asexual species-pairs found some overlapping genes, but these were close to the expectation by chance (Fig. 5B, for all levels see Supplemental Table 14). The majority of the DE genes also showed a significant interaction between species-pair and reproductive mode in the model used to identify convergently changing genes (whole-body = 69%, reproductive tract 66%, and legs = 81%), corroborating the finding that the vast majority of the DE genes are species-pair specific. Note the species-pair by reproductive mode interactions do not appear to be generated by one specific species-pair as generally genes DE between one species-pair were not DE between the other 4 species-pairs.

**Figure 5.**
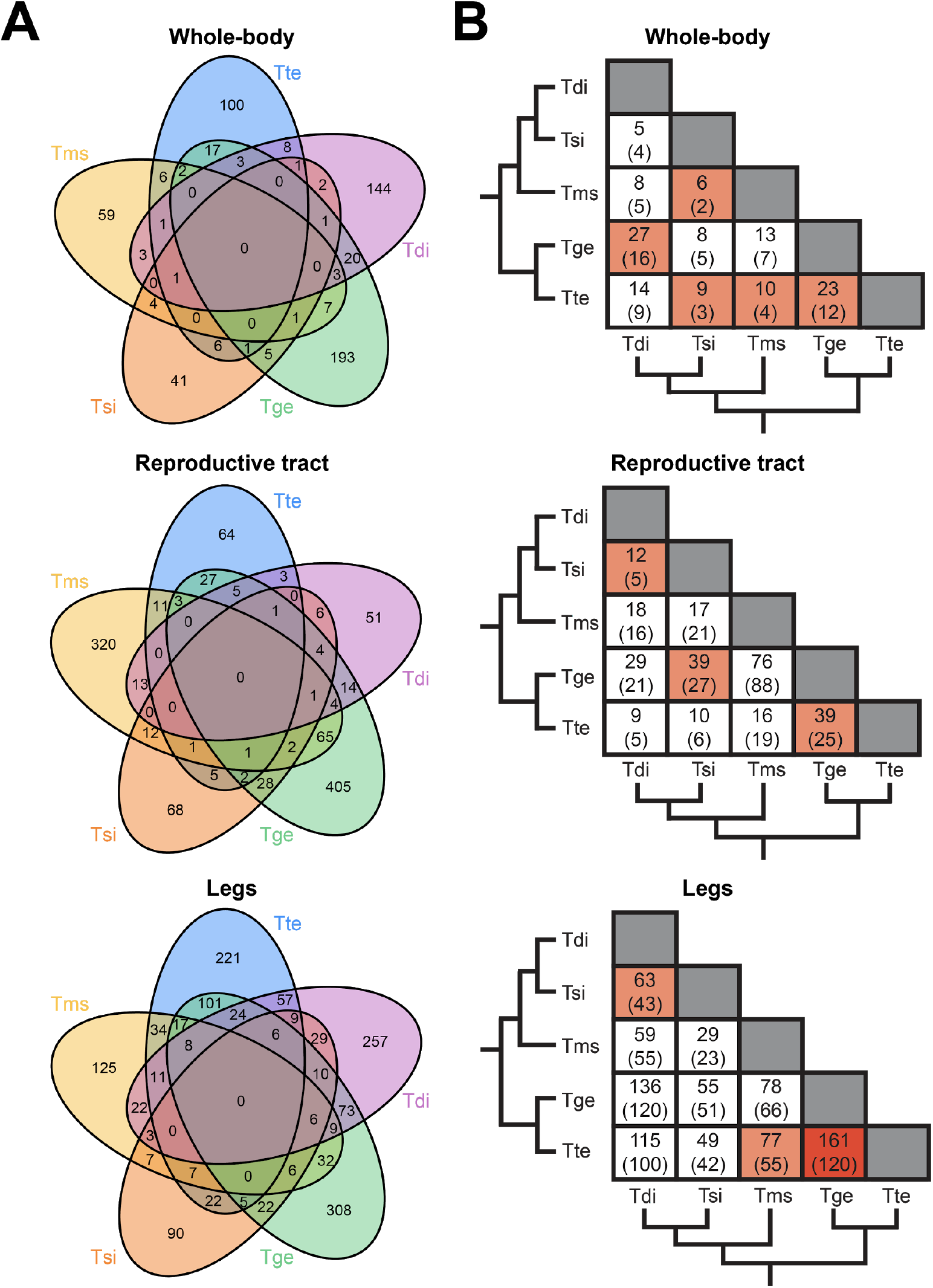
A. Venn-diagrams showing the number differentially expressed (DE) genes between sexual and asexual females that are shared among species-pairs for whole-body, reproductive tract, and legs for 10 species orthologs (FDR < 0.05). B. Matrices showing pairwise overlap of DE genes between sex-asex species with the number of genes expected by chance given in parentheses. Colours represent a significantly greater overlap than expected by chance (red, FDR < 0.001, orange < 0.05). The phylogeny shows the relationships between asexual species (from (2011)). Species-pair names are abbreviated as follows: Tte = *T. tahoe*, Tms = *T. monikensis*, Tdi = *T. douglasi*, Tsi = *T. shepardi*, and Tge = *T. genevievae*.

The DE genes of the different species pairs are not involved in convergent functional processes. Species-pair-specific genes were enriched for a number of GO terms (presented in Supplemental Table 15), however no GO terms were found to overlap between all pairs in any tissue (Supplemental Fig. 8). Examination of overlaps between pairs of sexual-asexual species-pairs found some overlapping GO terms, but these were close to the expectation by chance (Supplemental Table 16). This pattern remained even when a more liberal approach, whereby related GO-terms were considered as a unit, was applied (Supplemental Fig. 9A). This overall lack of overlap suggests that species-pair-specific genes are not involved in producing convergent phenotypes. Instead, species-pair-specific genes are the product either of lineage-specific selection (for instance, if different species use different mechanisms to achieve parthenogenesis) or of drift. These two processes are difficult to disentangle, but our results are more consistent with drift rather than with lineage-specific selection. Indeed, species-pair specific genes showed similar changes in gene expression across tissues, in contrast to the mainly tissue-specific changes uncovered for convergently changing genes. The overlap of species-pair specific genes between tissues was significantly greater than expected by chance (Table 1).

Finally, these results were reproduced when examining a much larger set of genes (genes with reciprocal-best-blast-hits between species-pairs, see above) as both genes DE between each species-pair, and their enriched GO terms, showed little overlap (Supplemental Figs 10 and 11, and Supplemental Tables 17 and 18).

## Discussion

Asexuality has convergently evolved numerous times across the tree of life, and a large body of research focuses on the reasons why sexual reproduction persists in the face of competition from asexual lineages. By contrast, the molecular underpinnings of transitions from sexual to asexual reproduction remain largely unknown (Neiman et al. 2014). In this study we examined gene expression changes associated with transitions to asexuality across five independently evolved asexual lineages, in whole-bodies, reproductive tracts and legs. The changes we observe provide, for the first time, insights into the convergent evolution of asexuality at the molecular level.

We found evidence for convergent changes in gene expression in all three tissues. Four lines of evidence suggest that these changes are a product of selection. Firstly, parallel changes across multiple independent transitions represent strong evidence of selection and thus are unlikely to be due to drift (Losos 2011; Natarajan et al. 2016). Secondly, convergent changes were primarily tissue-specific. This finding is consistent with selection, because expression changes due to drift are likely to be correlated across tissues (Blekhman et al. 2008; Liang et al. 2018). Indeed, the different functional roles of reproductive tracts and legs make it unlikely that selection would drive changes in the same genes in all tissues. Thirdly, expression changes of convergent genes inferred over the *Timema* phylogeny fit better with a model of adaptive change than with models of simple drift or of independent change in asexuals. Finally, the functional processes of convergently expressed genes mirror the changes observed at the phenotypic level, supporting the interpretation that these genes contribute to the convergent phenotypic changes observed between sexual and asexual females.

The overall amount of convergence is striking, particularly in the reproductive tract and legs with approximately 8% of genes showing a convergent shift in expression. Such a large amount of convergence suggests that the path from sexual reproduction to asexuality is strongly constrained, requiring changes to the same genes and biological processes in order to produce asexual phenotypes.

### Convergent changes in gene expression reveal the mechanisms underlying the production of asexual offspring

Asexuality is a complex adaptation that includes two major components: the ability to produce viable asexual offspring, and secondary adaptive changes that would not have been selected for in sexual species (e.g. the reduction of costly sexual traits). A key change necessary for the production of asexual offspring is the ability to produce unreduced eggs (Engelstädter 2008). Convergently expressed genes in the reproductive tract were enriched for changes in meiosis, and in particular meiotic spindles, which are key for the proper division of cells during meiosis. Mutations in meiotic spindles have been shown to result in unreduced meiotic products in *D. melanogaster*, and specifically in two genes (*bond* (Szafer-Glusman et al. 2008) and *pelo* (Eberhart and Wasserman 1995)) which show convergent changes in expression in asexual *Timema.* As such we suggest that these changes may underlie the non-reduction of eggs in asexual *Timema.* An alternative hypothesis is that since *Timema* reproduce parthenogenetically (and thus likely no longer recombine) changes in meiotic genes represent trait decay. Although possible, previous work has shown that, in fact, meiotic genes are not only retained in asexual lineages without damaging mutations, but often appear to be subject to selection for changes in expression, via duplication or differential upregulation of promoters (Srinivasan et al. 2010; Hanson et al. 2013; Raborn et al. 2016; Warren et al. 2018). Taken together with our results, we suggest that modifications to meiotic genes, specifically those that that disrupt meiotic cell division, are key in overcoming a major barrier to the evolution of asexuality: the production of unreduced eggs.

The production of unreduced eggs is not the only barrier to producing offspring asexually. In most species, sperm transfer essential components for the formation of a functioning centrosome (Schatten 1994; Manandhar et al. 2005). This paternal contribution represents a second key barrier in the evolution of parthenogenesis in many systems (Engelstädter 2008). However, in phasmids the centrosome is assembled without any contribution from sperm in both sexual and asexual species (Marescalchi et al. 2002). This may act as pre-adaptation for asexuality in stick insects and account, in part, for the large number of asexual stick insects.

A final barrier to asexual offspring production in many systems is egg activation. In many species mature oocytes are arrested at a specific stage (e.g. at metaphase II in mammals, and metaphase I in most insects), and must be activated by sperm to re-enter the cell-cycle (Stricker 1999; Engelstädter 2008; Nishiyama et al. 2010). In insects however, egg activation does not require sperm as activation is induced by the transit through the reproductive tract (Sartain and Wolfner 2013). Despite this, ovulation and egg-laying rates are strongly tied to mating (Eberhard 1996; Gillott 2003) meaning this signal must be modified in order for asexual insects to have normal levels of fecundity. In insects, the signal to a female that she has successfully mated is likely detected by sensory neurons in her reproductive tract (Yapici et al. 2008). Consistent with this, we find changes in gene expression linked to sensory neurons in the reproductive tract of asexual females, which may act to cue high levels of ovulation without mating. Alternatively, these changes may represent the decay of these neurons since they are no longer needed to detect mating events, or these changes may result from cessation of sexual conflict. Sensory neurons in the reproductive tract are known targets of substances in the male ejaculate (Heifetz and Wolfner 2004; Sakai et al. 2009; Heifetz et al. 2014) to induce the release of eggs and to reduce female receptivity (Gillott 2003). This manipulation is countered by female resistance adaptations which are likely costly, meaning that, following a transition to asexuality, there will be selection against them.

### Convergent changes in gene expression show evidence for the decay of female sexual traits

Sexual traits in asexual females are often observed to be reduced or lost (van der Kooi and Schwander 2014). For instance, in insects, females typically produce pheromones as a sexual cue to attract males (Greenfield 2002), and this cue has been repeatedly reduced or lost in several asexual species (see van der Kooi and Schwander (2014)), including *Timema* (Schwander, Crespi, et al. 2013). Such trait decay can be the result of either reduced purifying selection acting on traits that are now selectively neutral, or selection to reduce the cost of producing sexual traits. In asexual *Timema* reproductive decay has been primarily attributed to selection rather than reduced purifying selection, as reproductive trait decay in very young asexual lineages is as extensive as in old ones (Schwander, Crespi, et al. 2013).

Convergent gene expression changes underlying the decay of reproductive traits are mostly observed in *Timema* whole-bodies. In particular, we find enrichment of terms associated with sperm storage and sexual behaviour. Changes in the legs were less obviously associated with reproductive trait decay, however we do find changes in genes involved in cuticle development, pigment biosynthesis, sensory perception of touch, and changes in sexual behavior. These changes could represent the reproductive decay of both sexual cues (e.g. cuticular hydrocarbons and pigmentation which are both important for mate choice in insects (reviewed in Hunt and Sakaluk (2014)), and their detection (via sensory receptors on the leg (reviewed in Dahanukar et al. (2005)). In addition, we also find changes in genes associated with sex determination in the soma, including *sex-lethal*, a master-feminizing switch in *Drosophila* (Cline et al. 2010) which may have a major influence the development of many sexual traits in the legs.

Although we focus on expression, it is possible that the decay of sexual traits is also evident at the sequence level. By examining the coding regions of genes, we found evidence for reduced purifying selection acting on the sequence of genes showing convergent expression changes in the whole-body. This suggests, that in some cases, the reduction of sexual traits may be accomplished by both expression and sequence changes, which potentially act interactively to produce a phenotypic change.

Unexpectedly, we also find changes to immune function in the legs and whole-body, the majority of which show down-regulation in asexual females. A possible explanation for this is that asexual females are likely to face a reduced number of immune challenges compared to sexual females due to the elimination of sexually transmitted diseases, the costs of which can be considerable, even shaping the evolution of many aspects of an organism’s life history, such as mate choice, mating rate, and sexual signal investment (Kokko et al. 2002; Knell and Webberley 2004). As such we suggest asexual females may be reducing the allocation of resources to immune function due to the absence sexually transmitted diseases. This effect may be particularly strong in solitary species such as *Timema*, where the majority of socially transmitted diseases come from sexual interactions.

### Species-pair specific changes

In addition to convergent changes, we also identified many species-pair specific gene expression changes. In contrast to convergent genes, species-pair specific genes showed common shifts in expression across tissues, and inconsistent associations with functional processes between species-pairs, that were largely unrelated to asexual phenotypes. Taken together, these results suggest that the majority of changes we observe from a single sex-asex species-pair comparison are due to drift rather than selection. Our findings thus highlight the problem of drawing inferences on the causes or consequences of asexuality from the examination of only a single transition to asexuality, whereas examining several transitions allows us to disentangle adaptive changes and those due to drift.

Overall, we find evidence for a striking number of convergent changes across five transitions to asexuality. Previous studies that have examined convergent expression changes across the genome have found little evidence of convergence at the gene level (<1%) (Rittschof et al. 2014; Berens et al. 2015), underlining the surprisingly large amount of convergent gene expression (up to 8%) we find here. The amount of molecular convergence to expect, however, is dependent on several factors including the complexity of the phenotype, and the size of the mutational target (Martin and Orgogozo 2013). For example, we find that a key change required for asexual reproduction, the production of unreduced eggs, likely requires changes to meiotic spindle regulation. The pathways that govern meiotic spindle regulation are relatively small in number (Bennabi et al. 2016), meaning that only a small minority of genes are likely able to confer the relevant changes, making the chance of molecular convergence for this trait relatively high. By contrast, the observed reduction of sexual traits could be produced by changes to numerous genes and pathways (i.e. there is a large mutational target) making convergent molecular changes for these traits less likely. Despite this, our and previous studies examining trait loss have also demonstrated a high amount of convergence (Whittall et al. 2006; Gross et al. 2009; Partha et al. 2017; Sackton et al. 2018), implying that certain genes have a disproportionate role in not only the convergent evolution of novel phenotypes, but also in their convergent loss (Martin and Orgogozo 2013; Stern 2013).

## Methods

### Samples

Females for whole-body samples were collected from the field as juveniles in spring 2013. All individuals were then raised in common garden conditions (23°C, 12h:12h, 60% humidity, fed with *Ceanothus* cuttings) until eight days following their final molt. Prior to RNA extraction, individuals were fed with artificial medium for two days to avoid RNA contamination with gut content and then frozen at −80°C. Individuals used for tissue-specific samples were collected in spring 2014 as juveniles and raised in the same common-garden conditions as whole-body samples. For leg samples three legs were used from each individual (one foreleg, one midleg, and one hindleg). Reproductive tracts were dissected to consist of ovaries, oviducts and spermatheca. Note the same individuals were used for leg and reproductive tract samples. Collection locations for all samples are given in Supplemental Table 19.

### RNA extraction and sequencing

The three biological replicates per species and tissue consisted of 1-9 individuals per replicate, which were combined prior to RNA extraction (207 individuals in 90 replicates in total; see Supplemental Table 19). RNA extraction was performed by freezing samples in liquid nitrogen followed by addition of Trizol (Life Technologies) before being homogenized using mechanical beads (Sigmund Lindner). Chloroform and ethanol were then added to the samples and the aqueous layer transferred to RNeasy MinElute Columns (Qiagen). RNA extraction was then completed using an RNeasy Mini Kit following the manufacturer’s instructions. RNA quantity and quality was measured using NanoDrop (Thermo Scientific) and Bioanalyzer (Agilent). Strand-specific library preparation (1 library per replicate) and single-end sequencing (100 bp, HiSeq2000) were performed at the Lausanne Genomic Technologies Facility.

The 90 libraries produced a total of just over 3 billion single-end reads. Four whole-body and six tissue-specific libraries produced significantly more reads than the average for the other samples. To reduce any influence of this on downstream analyses, these libraries were sampled down to approximately the average number of reads for whole-body or tissue-specific libraries respectively using seqtk (https://github.com/lh3/seqtk Version: 1.2-r94).

### Transcriptome references

*De novo* reference transcriptome assemblies for each species were generated previously (Bast et al. 2018). For our analyses we used the 3010 one-to-one orthologs present in all 10 *Timema* species as identified by Bast et al. (2018). Identified ortholog sequences varied in length among different species. Since length variation might influence estimates of gene expression, we aligned orthologous sequences using PRANK (v.100802, default options) (Löytynoja and Goldman 2005) and trimmed them using alignment_trimmer.py (Parker 2016) to remove overhanging gaps at the ends of the alignments. If the alignment contained a gap of greater than 3 bases then sequence preceding or following the alignment gap (whichever was shortest) was discarded. Three genes were discarded at this stage as the trimmed length of sequence was <300 bp. These trimmed sequences were then used as reference transcriptomes for read mapping. Note that genes with significant Blast hits to rRNA sequences were removed from the transcriptome references prior to mapping.

### Read trimming and mapping

Raw reads were trimmed before mapping. Firstly CutAdapt (Martin 2011) was used to trim adapter sequences from the reads. Reads were then quality trimmed using Trimmomatic v 0.36 (Bolger et al. 2014): first clipping leading or trailing bases with a phred score of <10 from the read, before using a sliding window from the 5’ end to clip the read if 4 consecutive bases had an average phred score of <20. Following quality trimming any reads <80 bp in length were discarded. Quality-trimmed reads from each library were then mapped separately to the reference transcriptome using Kallisto (v. 0.43.1) (Bray et al. 2016) with the following options -l 210 -s 25 --bias --rf-stranded for whole-body samples and -l 370 -s 25 --bias --rf-stranded for tissue specific samples (the -l option was different for whole-body and tissue specific samples as the fragment length for these libraries was different).

### Differential expression analysis

Expression analyses were performed using the Bioconductor package EdgeR (v. 3.18.1) (Robinson et al. 2010) in R (v. 3.4.1) (R Core Team 2017). Analyses were done separately for each tissue. Genes with counts per million less than 0.5 in 2 or more libraries per species were excluded from expression analyses. Normalization factors for each library were computed using the TMM method in EdgeR. To estimate dispersion we then fit a generalized linear model (GLM) with negative binomial distribution with the terms species-pair, reproductive mode and their interaction. We used a GLM likelihood ratio test to determine significance of model terms for each gene by comparing appropriate model contrasts. P-values were corrected for multiple tests using Benjamini and Hochberg’s algorithm (Benjamini and Hochberg 1995), with statistical significance set to 5%. Using this approach, we classified genes as convergently differentially expressed when there was a significant effect of reproductive mode (FDR < 0.05) but no interaction effect of species-pair by reproductive mode (FDR > 0.05). Differentially expressed genes within each species-pair were identified using pairwise contrasts between each sexual and asexual pair.

To determine if genes DE within each species-pair and tissue show greater than expected number of overlapping genes we used the SuperExactTest package (v. 0.99.4) (Wang et al. 2015) in R which calculates the probability of multi-set intersections. When examining multiple intersections p-values were multiple test corrected using Benjamini and Hochberg’s algorithm implemented in R.

To test if the observed number of convergent genes was significantly greater than expected by chance we performed a permutation test where, for the read counts of each gene, we randomly switched the assignment of reproductive mode (sexual or asexual) within a species-pair. Note that all biological replicates from a particular group were always assigned to the same reproductive mode (i.e. In the event of a switch, all sexual replicates were assigned as asexual, and vice versa). This process was repeated to produce 10,000 permuted data sets, which were then run through the gene expression pipeline described above to generate a distribution of the number of convergent genes we expect to find by chance.

### Ornstein-Uhlenbeck based models

Ornstein-Uhlenbeck (OU) based models were fit using the R package ouch (v. 2.11-1) (King and Butler 2009). We fit three models to each convergently expressed gene: a drift-model where changes are modelled as Brownian motion, a two-state adaptive-optima model where asexual species had a different adaptive-optima than sexual species, and a multi-asexual optima model where sexual species and each asexual species had different optima (Supplementary Fig. 12). Initial values of sqrt.alpha and sigma for the adaptive-optima models were set to 1. Log-likelihood ratio tests were used to determine if the two-state adaptive-optima model was a significantly better fit to observed expression values than the drift-model, and if the multi-asexual optima model was a significantly better fit to observed expression values than the two-state adaptive-optima model. P-values were corrected for multiple tests using Benjamini and Hochberg’s algorithm (Benjamini and Hochberg 1995), with statistical significance set to 5%. Mean observed expression values (log_2_ counts per million) for each species and gene were calculated using EdgeR (see above). We used RAxML (v. 8.2.8, options: -p 12345 -m GTRCAT -T 6) (Stamatakis 2014) to produce a maximum likelihood tree for use in ouch using the concatenated one-to-one ortholog alignments produced in Bast et al. (2018).

### GO term analysis

Genes were functionally annotated using Blast2GO (version 4.1.9) (Götz et al. 2008) as follows: sequences from each sexual species were compared with BlastX to either NCBI’s nr-arthropod or *Drosophila melanogaster* (drosoph) databases, keeping the top 20 hits with e-values <1 x 10^−3^. Interproscan (default settings within Blast2GO) was then run for each sequence, and the results merged with the blast results to obtain GO terms. This produced two sets of functional annotations, one derived from all arthropods and one specifically from *Drosophila melanogaster*. The *D. melanogaster* GO term annotation generated around four times more annotations per sequence than NCBI’s nr-arthropod database. We therefore conducted all subsequent analyses using the GO terms derived from *D. melanogaster* but note that results using the annotations from all arthropods were qualitatively the same (see Supplementary Fig. 9B).

We conducted gene set enrichment analyses using the R package TopGO (v. 2.28.0) (Alexa and Rahnenfuhrer 2016) using the elim algorithm to account for the GO topology. Gene set enrichment analyses identify enriched GO terms in a threshold-free way, by finding GO-terms that are overrepresented at the top of a ranked list of genes. For comparisons within a species-pair, genes were ranked by FDR; to identify enrichment of convergent genes, genes were ranked by FDR value for reproductive mode, with the FDR value for genes that showed a significant lineage by reproductive mode set to 1. GO terms were considered to be significantly enriched when p < 0.05. Enriched GO terms were then semantically clustered using ReviGO (Supek et al. 2011) to aid interpretation.

The significance of overlapping GO terms was determined using SuperExactTest as described above. The hierarchical nature of GO terms generates a bias towards finding a significant amount of overlap, since enrichment terms are non-independent. It is however possible that the complexity of the GO term hierarchy could lead to convergent functional processes being overlooked. For instance if a GO term is enriched in one comparison, but its parent term is enriched in another comparison, then there would be no apparent overlap. To address this, we also looked at the amount of ‘linked overlap’ of GO terms, whereby significant GO terms were first clustered together based on parent or child terms.

For the GO term enrichment analyses of convergently differentially expressed genes we used only the annotation from *T. bartmani* as it had the most number of sequences annotated. Annotations to each of the other species were very similar to those from *T. bartmani,* with 80% of annotations being identical across all 5 species annotations. The remaining 20% of sequences were typically characterized by an additional term in one or more of the species. For comparisons within a lineage we used the annotation of the sexual species in that lineage. Although the annotations are very similar across all ten species the small differences in annotation could create differences in the amount of overlap observed between contrasts (e.g. if a term is annotated to an ortholog in one annotation but not another). To examine this, we repeated the analysis using only annotations from *T. bartmani*. This produced a virtually identical result (Supplemental Fig. 9C) as when using the species-pair specific annotations.

### Polymorphism and divergence

To test for differences in the rate of evolutionary divergence between gene categories, we used dN/dS ratios for each of the one-to-one orthologs from Bast et al. (2018). Briefly, Bast et al. (2018) aligned each of the one-to-one orthologs with M-Coffee (Wallace et al. 2006) and trimmed them with Gblocks (Talavera and Castresana 2007). These alignments were then used as input for codeml of the PAML package (Yang 2007) to generate maximum likelihood estimates of dN/dS for each terminal branch in the phylogeny (using the “free model”). To obtain an estimate for pN/pS for each ortholog, reads from the whole-body libraries (based on a pool of three individuals) for each asexual species were mapped to the reference using RSEM/bowtie2 with default parameters and fragment length mean = 200 fragment length sd = 100 (Li and Dewey 2011; Langmead and Salzberg 2012). Samtools v1.2 was then used to create an mpileup file, which was filtered with VarScan v2.3.2 (minimum coverage = 20, minor allele frequency = 10%, and minimum average phred quality = 20) to obtain SNPs. To identify nonsynonymous and synonymous segregating polymorphisms we identified the n-fold degenerate positions following Li et al. (1985) from which pN, pS and (pN/pS) could be calculated per gene. Comparison of mean pN/pS and dN/dS between convergent and non-convergent (background) genes was conducted using a permutation t-test (number of permutations = 10000) in R.

### Cross-species mapping

All of the above analyses used only the one-to-one orthologs. To examine a larger fraction of the transcriptome we produced species-pair references by using a reciprocal blast between the assemblies of sexual-asexual sister species (*T. bartmani* - *T. tahoe, T. cristinae - T. monikensis*, *T. poppensis - T. douglasi*, *T. californicum - T. Shepardi*, and *T. podura - T. genevievae*) (blastN, minimum e-val = 0.00001, minimum query coverage = 30%). Prior to this step potential contaminants were filtered from these by blasting transcripts to local versions of the nt (using blastN, default options except task = blastn, max_target_seqs = 10) and nr (using blastX, default options except, max_target_seqs = 10) databases (downloaded: 07/08/2016) using NCBI’s blast client (v. 2.2.30+). Blast hits with an e-value > 0.0000001 were discarded. The remaining blast hits were used to assign a phylum to sequences if >=50% of Blast hits came from one phylum (in the event of a tie, the taxa with the highest e-value was used as a tiebreaker). Transcripts that were assigned to a non-arthropoda phylum were discarded (note that transcripts with no Blast hits or that blasted to mixed phyla were retained). This filtering removed between 4-8% of transcripts (see Supplemental Table 10). Reads of each species were then mapped to each species-pair references in the same way as for the 1-to-1 orthologs. Differential expression analyses and GO-term enrichment analyses were then repeated as described above.

## Data

Raw reads have been deposited in the SRA. Accession codes are given in Supplementary Table 19. Scripts for the analyses in this paper are available at: https://github.com/DarrenJParker/Timema_convergent_gene_expression.

## Acknowledgments

This study was supported by Swiss FNS grants PP00P3_170627, PP00P3_139013, and CRSII3_160723. We would like to thank Chloé Larose and Bart Zijlstra for their assistance in the field.

## Author contributions

T.S., D.J.P. and K.J. designed the study. Z.D., K.J. and TS collected samples, performed dissections and molecular work. D.J.P. analyzed the data with input from J.B., T.S. and M.R.R. D.J.P. and T.S. wrote the manuscript with input from all authors.

